# Identification of the yeast mannoprotein gene *HZY1* as a key genetic determinant for yeast-derived haze in beer

**DOI:** 10.1101/2023.07.10.548400

**Authors:** Keith Lacy, Rita Mormando, Jeremy R. Smith, Patrick A. Gibney, Lance M. Shaner, Laura T. Burns

**Affiliations:** Omega Yeast Labs, Chicago, IL, U.S.A.; Department of Food Science, Cornell University, Ithaca, NY, U.S.A.

**Keywords:** haze, colloidal haze, turbidity, dry hop, hazy IPA, mannoprotein, intragenic repeats, beer

## Abstract

With the sustained popularity of hazy IPAs, brewers have explored multiple approaches to maximizing stable haze that will remain in suspension throughout the shelf life of the beer. Our recent investigations into yeast-dependent haze have uncovered specific brewing yeast strains that promote the formation of haze in heavily dry-hopped beer styles. These brewing strains have been termed “haze-positive” and furthermore, the timing of dry hop additions has been found to be another key factor in producing this stable haze. Classical genetics have identified YIL169C (herein referred to as *HZY1*) as both necessary and sufficient for the haze-positive phenotype in the yeast strain most widely used for Hazy IPAs. *HZY1* encodes a candidate glycoprotein and our recent findings suggest it is localized to the cell wall through a GPI anchor. Surprisingly, using long-read sequencing data we uncovered extensive genetic variation in *HZY1* across brewing strains. The haze-positive phenotype correlates with an expansion in the N-terminal serine-rich region. We propose that the Hzy1 glycoprotein is a critical component to yeast-dependent colloidal haze and the genetic variation in this locus contributes the range of haze phenotypes observed across industrial brewing strains.

## Introduction

Hazy IPAs (also referred to as Juicy IPAs, New England IPA, East Coast IPA, NEIPA) have grown rapidly in popularity over the past decade, now representing at least 10 percent of the craft beer market. With the fast growth in hazy beer styles, brewers have faced challenges in producing stable haze that persists in the beer throughout the product shelf life. Prior beer haze research in lagers and wheat beers indicated that malt and hops were the major drivers of haze, with a mechanism involving protein-polyphenol interactions ^1-3^. Many brewers early on had the misconception that non-flocculent yeast was responsible for producing haze, but this approach provided a very short-term haze and became problematic as the yeast settled out after packaging into kegs and cans. Through trial and error, one yeast strain, OYL-011 (trade names include British V, London Ale III, Foggy London, and Juice), became widely popular for Hazy IPAs due to its fruity fermentation profile, lower attenuation, and ability to produce a reliable, stable haze. As this growing category of Hazy IPAs dominated the market, we set out to better understand the genetics behind yeasts’ ability to promote stable haze in dry-hopped beer styles.

There has been a strong link between haze and yeast mannoproteins in beer and wine. Yeast mannoproteins are heavily glycosylated cell wall-derived proteins and are responsible for flocculation, biofilm and cell sensing. In beer, stressed yeast has been proposed to result in increased turbidity. Early studies examined the impact of agitation and shear stress and found the release of mannoproteins led to increased haze and difficulties in subsequent beer filtration. A recent investigation into sporadic haze across a brewery’s IPA production suggested that yeast mannoproteins and poor yeast management were key factors leading to increased haze ^4^. Counter to the haze promoting role for yeast mannoproteins in beer, mannoproteins have been shown to prevent the formation of haze in wine. There is a strong negative correlation of glycoprotein to haze in wine and the overexpression of some mannoproteins results in decreased haze formation ^5-8^. Non-*Saccharomyces* strains have been found to release polysaccharides in the form of glycoproteins as well, promoting desirable mouthfeel while also reducing haze. Thus, it is important to note that *Saccharomyces cerevisiae* mannoproteins may have different effects on haze in different fermentation matrices and the role for specific mannoproteins in beer haze is largely still unknown. Here, we established an assay for phenotyping yeast-derived haze and leveraged classic backcrossing and modern whole genome sequencing to identify the genetic component for haze in OYL-011. This approach has uncovered complex genetic variation in the *HZY1* gene and provides the first established role for this novel mannoprotein in promoting stable colloidal haze in beer.

## Methods

### Assay for Yeast-Dependent Haze

Small scale fermentations of brewer’s wort prepared from barley malt were inoculated with each candidate yeast strain to 10 million cells/ml. Separate flask fermentations were dry hopped with 8 g/L of T90 pellet hops for each day following inoculation until day 7. After a total of 14 days of fermentation, the beer samples were centrifuged at 5000 rpm for 5 minutes to remove all yeast cells and particulate. The resulting clarified beer samples were measured for haze in an Anton Paar HazeQC turbidity meter.

### Yeast Strains and Genetic Backcrossing

Genetic manipulations including crosses, sporulation and tetrad analysis were carried out using standard procedures ^9^. Yeast strains used in this study can be found in the strain table (Supplemental Table S1). For genetic backcrossing experiment, the OYL-011, a haze-positive tetraploid heterozygous industrial brewing strain, was backcrossed to a haze-neutral homozygous diploid wine strain. At each backcross, the resulting haze-positive isolates were used for subsequent backcrosses until the seventh backcross in which the resulting isolates were approximately 99.2% identical to the wine strain parent.

### Variant Calling and Variant Distribution Mapping

Full genome sequences with >1 Gbp of sequencing data were obtained for the parent strains and four isolates from the seventh backcross (BC7-A, BC7-B = haze-positive and BC7-C, BC7-D = haze-neutral). Quality trimming and adapter clipping was performed using Trimmomatic v0.29 with default parameters ^10^. The trimmed raw reads were mapped to the *S. cerevisiae* reference genome (S288C) using Minimap2 v2.24 ^11^. From the resulting alignments, variants were called using Freebayes v0.9.21 ^12^. A custom bash script was used to parse through the resulting VCF files of each sample. A custom Python script was used to generate variant distribution plots corresponding S288C genome coordinates for each strain using Seaborn v0.12.2 and Matplotlib v3.7.0 Python packages. The position of each variant call within the four isolates was compared to the position of each variant call within the parent wine strain. A genotype ratio of the BC7 isolate to parent wine strain was plotted with a 50 bp sliding window with a step size of 5 along the S288C genome coordinates.

### Cloning of *HZY1* Alleles

The haze-positive isolate from BC7 (OYR-329) was dissected and tetrads were PCR confirmed for the OYL-011 and wine strain alleles of *HZY1*. Each allele was PCR amplified and subcloned into a shuttling vector with AMP and HYG-B drug-resistant cassettes. The resulting vectors were sequenced with oxford nanopore long-read sequencing. The OYL-011 *HZY1* allele was unstable and exhibited frequent loss of the N-terminal and C-terminal repeat motifs and thus the PCR product was also sequenced with oxford nanopore long-read sequencing.

### Violin Plots of *HZY1* N-term and C-term lengths

Full long-read genome sequences were obtained for the Omega Yeast Labs collection using Oxford Nanopore sequencing. Three sequence tags were designed in the highest conserved regions surrounding and within the *HZY1* gene (Supplemental Table S2). All reads were mapped to the tag1 sequence using Minimap2 v2.24 ^11^. The resulting paf file was converted to a bed file resulting in subsequence fragments starting/ending with the sequence tag using a custom python script. The subsequences were then extracted using BEDTools v2.30.0 ^13^. Using the same methods, the tag1 containing reads were mapped to the tag2 sequence and the resulting sequences trimmed to tag1 and tag2 were combined into one fasta file. The length of all the trimmed reads for each sample were obtained using the sequence-stats v1.1 bash package. Violin plots were generated to show the distribution of N-term lengths using the Ggplot2 v3.4.2 package in R ^14,15^. The C-term plots were made following the same methodology as listed above with the exception that the reads were first mapped to tag2 and then the tag3 sequence.

### *HZY1* Disruption

Plasmids and oligonucleotides can be found in Supplemental Table S1. *HZY1* was disrupted in OYL-004, OYL-011, OYL-009 and OYL-077 using CRISPR/Cas9 gene editing. A plasmid containing the CRISPR/cas9 enzyme, *HZY1* targeting sgRNA, G418 drug-resistant cassette (pOY108) and an ssDNA oligo repair template with homology to the 5’UTR and 3’UTR (oligo 256) was transformed using standard Li/Ac transformation protocol ^16^. Selection with G418 was used to screen for pOY108 transformants. Subsequently, G418+ transformants were screened to confirm loss of endogenous *HZY1* (primers 229/257) and the resulting scar after *HZY1* deletion (primers 257/258). The corresponding *hzy1Δ* isolates were outgrown and G418-isolates were selected for further experiments.

### Brewing Trial and Tetrad Test

An IPA wort was prepared with 85% 2-row base malt and 15% Munich malt to 16.9 plato. Hot side hop additions included 1 g/L Mosaic at 10 minutes remaining in the boil and 2 g/L of Citra at the beginning of a 15-minute whirlpool. The wort was chilled to 68°F, aerated with oxygen and transferred to two fermentation vessels. OYL-011 or OYL-011 *hzy1Δ* yeast strains were pitched at 10 million cells/ml and fermentations were maintained at 70°F. On day 7 of fermentation, the beers were dry hopped with 16 g/L of Citra. The fermentations were complete on day 14 and were cooled to 32°F for 4 days before transferring to serving vessels to carbonate. Tetrad sensory analysis was performed with a panel of eleven tasters. The beers were served blind in opaque cups and covered to ensure tasters were unable to see differences in the turbidity.

## Results

To begin investigating the elusive role for specific yeast strains in promoting haze in beer, we developed an assay for phenotyping haze. We started by testing the popular Hazy IPA strain, OYL-011, and the OYL-004 control strain commonly used for non-hazy American craft styles. Fermentations were performed in 500 ml flasks and dry-hop additions were added during the fermentation to promote the formation of haze. After fermentation was complete (14 days), the samples were collected, centrifuged and measured for haze. The late fermentation dry hop additions between day four and day seven showed the greatest difference in haze measurements in the OYL-011 and OYL-004 strains (Figure 1A and 1B, respectively). The day seven dry hop was used to phenotype a collection of brewing strains (Figure 1C). A range of haze phenotypes was observed and an arbitrary cut off of 200 NTUs was used to define haze-positive (>200 NTUs) and haze-neutral (<200 NTUs) strains. Traditional English and American ale strains were among the most haze-positive strains, whereas Belgian ale strains and German lager strains were the most haze-neutral.

**Figure 1.**
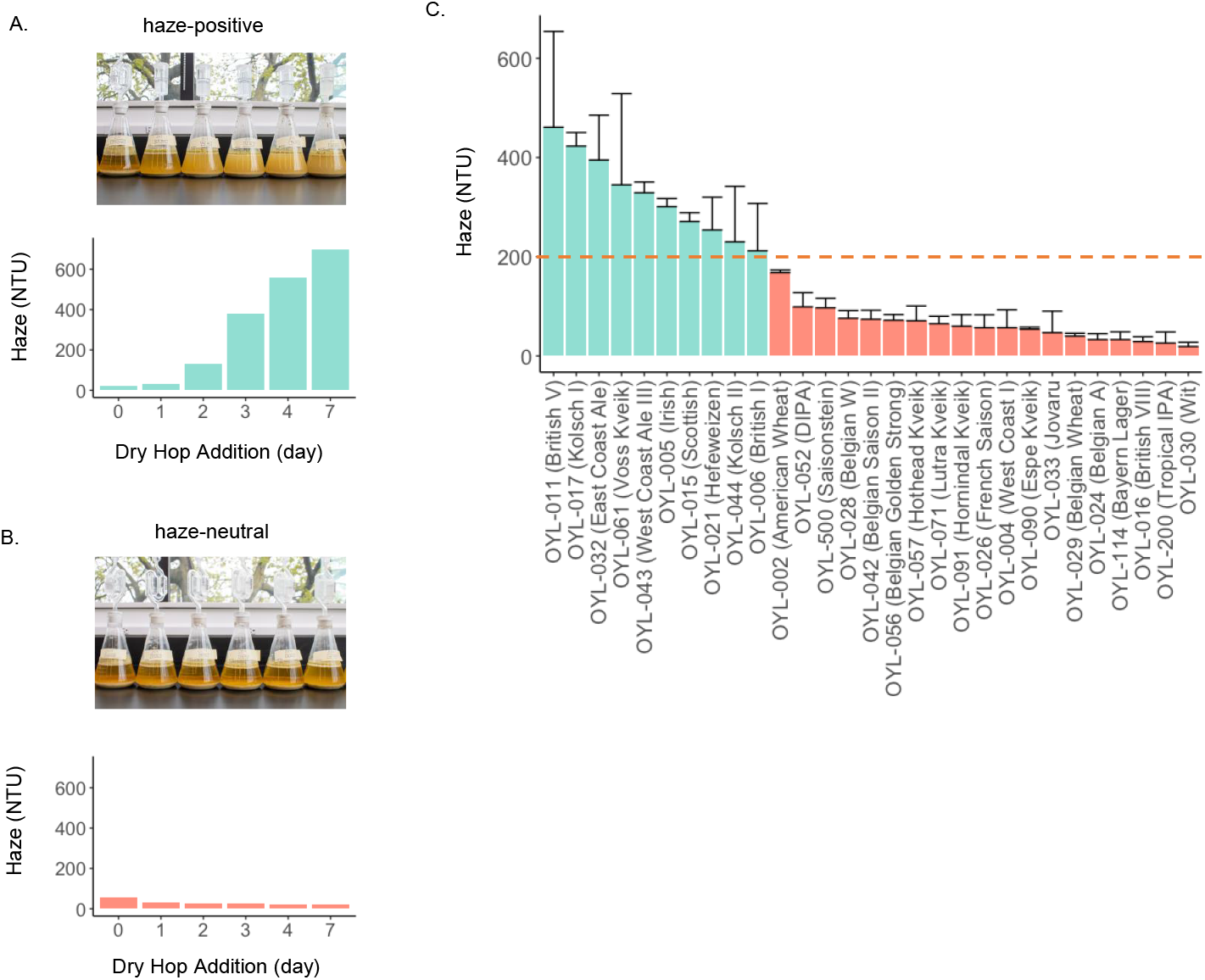
Characterization of the haze phenotype. (A) Image and haze measurements documenting the haze-positive phenotype of OYL-011. (B) Image and haze measurements documenting the haze-neutral phenotype of OYL-004. (C) Measurements of haze with day seven dry-hop addition in a collection of brewing strains. The average of a minimum of three experimental replicates are plotted for each strain with error bars representing standard deviation. Red dashed line at 200 NTUs indicates cutoff to define haze-positive and haze-neutral phenotype.

With an established haze phenotyping assay, we set out to determine the genetics of haze. We began by crossing the haze-positive OYL-011 to a haze-neutral wine strain (Figure 2A). We chose the Maxithiol wine strain for these crossing experiments because it was previously established to be homozygous diploid and therefore could be further used for backcrossing experiments. From the first cross, three hybrid strains were obtained, two haze-positive and one that was haze-neutral indicating the possibility of the haze-positive phenotype linking to a dominant Mendelian trait. We selected one of the haze-positive isolates and continued through seven successive backcrosses (Figure 2B). At backcross 7 (BC7) we obtained two haze-positive (BC7-A, BC7-B) and haze-neutral isolates (BC7-C, BC7-D) that were 99.2% isogenic to the wine strain parent (Figure 2C). We used Illumina sequencing to obtain >1 Gbp of whole genome sequencing data for the parent strains along with the BC7-A, BC7-B, BC7-C and BC7-D isolates. Variant calling was performed for the BC7 isolates and the wine strain against the reference genome S288C. Variant frequencies were plotted by the corresponding genome coordinates as a ratio between the BC7 isolates to the wine strain (Supplemental Figure S1). One region on the left arm of chromosome IX (0-30,000 bp) contained the candidate haze locus, as it was the only region that contained variants unique to the BC7-A and BC7-B haze-positive isolates and not the wine strain parent or BC7-C and BC7-D haze-neutral isolates (Figure 2D).

**Figure 2.**
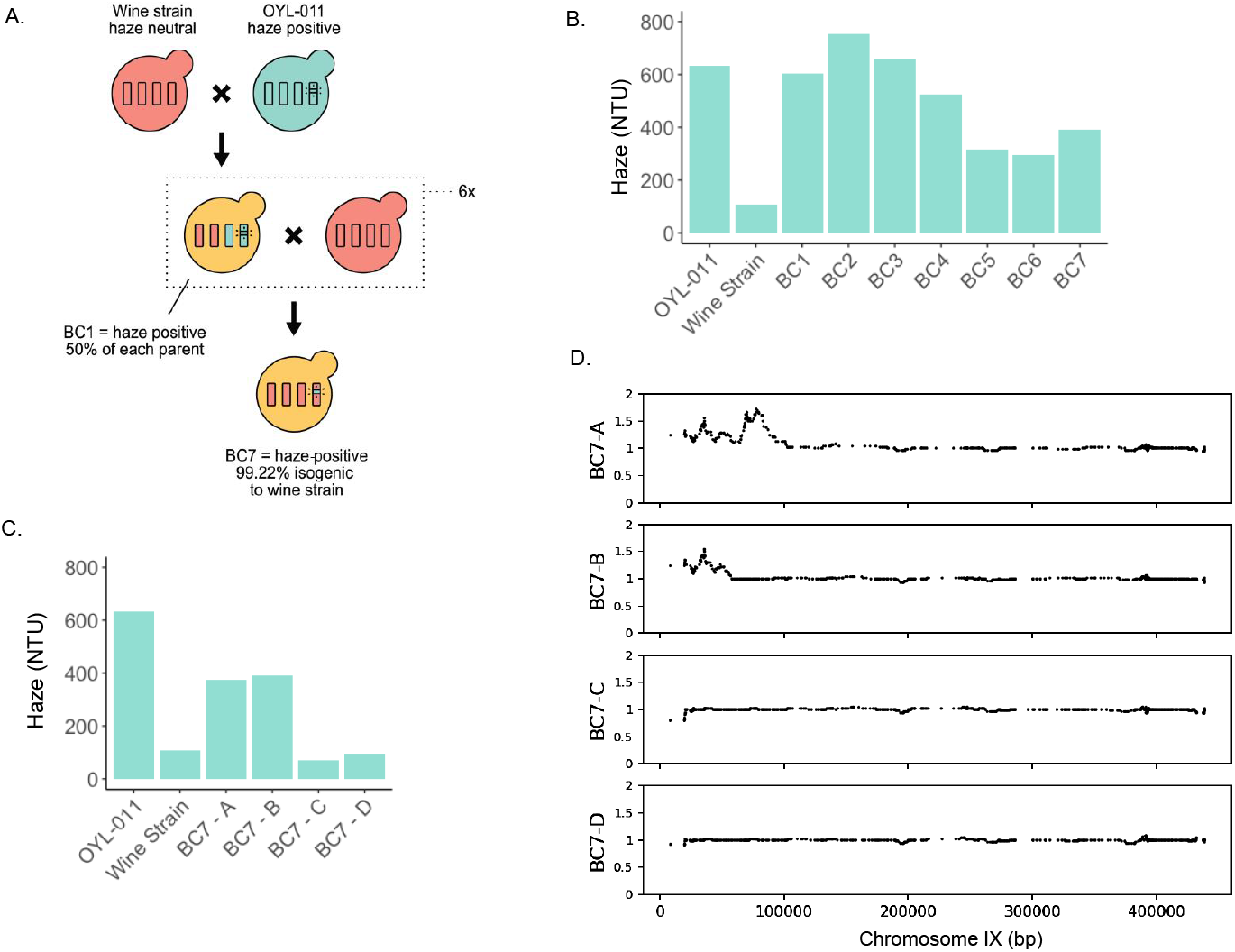
Backcrossing OYL-011 and identification of candidate haze locus in left telomeric region of Chr IX. (A) Schematic representation of the OYL-011 and wine strain backcrossing. (B) Haze measurements of haze-positive isolates from each backcross. (C) Haze measurements of the parent strains and the two haze-positive and two haze-neutral BC7 isolates (D) Variants specific to the two haze-positive BC7 isolates map to a candidate haze locus on the left arm of Chr IX. The ratio of variants found in the BC7 isolates relative to the parent wine strain (y-axis) are plotted with a sliding window of 50 bp along the Chr IX coordinates of the reference genome S288C (x-axis).

To further narrow down this region of chr IX to a candidate haze gene, we mapped the short read sequencing data of the parent strains and BC7 isolates to the S288C reference genome. The region between 23,000-26,000 bp showed a several-fold increase in coverage in the OYL-011 parent and BC7-A and BC7-B isolates, suggesting two potential repeat expansions within YIL169C at 25,400 bp and 23,800 bp corresponding to the N-terminus and C-terminus (Figure 3A-B). Since the BC7-A and BC7-B isolates were heterozygous for YIL169C, we sporulated BC7-A to obtain meiotic segregants that were homozygous diploid and either haze-positive or haze-neutral. We then assayed the meiotic segregants for haze. Primers were designed to amplify the N-terminus of YIL169C. Two products of distinct size were identified, with the “long” allele corresponding to the haze-positive phenotype (Figure 3C). Furthermore, PCR products of the full-length OYL-011 “long” and wine strain “short” alleles of YIL169C were directly sequenced using nanopore sequencing and aligned to the YIL169C allele of S288C, confirming the haze-positive OYL-011 allele contains expansions in both the N-terminal and C-terminal regions (Figure 3D). From the amino acid sequence, we noted that the N-terminus is heavily enriched in serine (OYL-011 “long” >55%) and the C-terminus in serine and threonine (OYL-011 “long” >45%). Repeat motifs rich in serine and threonine have been reported in other cell wall proteins ^17^. When further examining the YIL169C alleles, we identified two repeat motifs, a 14 aa motif in the N-terminus and a 36 aa motif in the C-terminus (Figure 3E). The N-terminal repeat was expanded 53 times in the OYL-011 strain, but was only found 15 times in the S288C lab strain and 7 times in the wine strain. The C-terminal repeat was expanded 17 times in the OYL-011 strain, while only identified once in both the S288C lab strain and wine strain. Another unanticipated finding was a predicted GPI anchor present in the wine parent strain and OYL-011, but not in the S288C lab strain. The loss of the GPI anchor in S288C suggests that the name for YIL169C, *CSS1* for “condition-specific secreted,” named as a part of a large-scale localization screen, may not be accurate. Therefore, we have renamed YIL169C, *HZY1*, and herein provide the first evidence that *HZY1* is necessary for promoting stable colloidal haze in beer.

**Figure 3.**
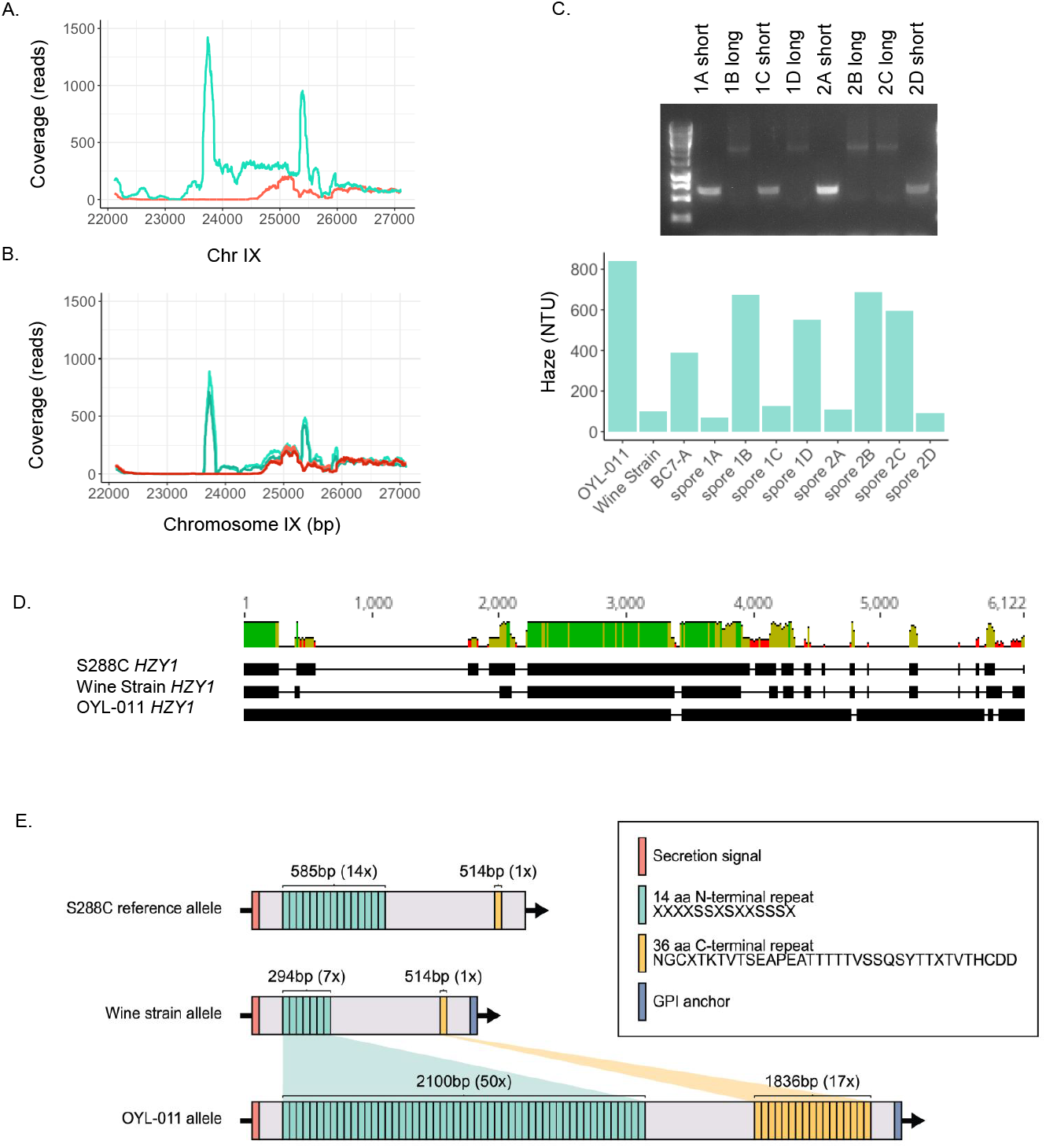
Large intergenic repeat expansions in the OYL-011 *HZY1* allele are associated with haze. (A) Coverage plot using short reads from Illumina whole genome sequencing from the parent wine strain (red line) and the parent OYL-011 strain (blue line) mapped to S288C reference genome. Regions within the N-term and C-term show increased coverage in the OYL-011 strain indicating potential repeat expansions in regions of *HZY1*. (B) Coverage plot using short reads from Illumina whole genome sequencing for the BC7-A and BC7-B isolates (light green, dark green) and BC7-C and BC7-D (light red, dark red). BC7-A and BC7-B isolates also exhibit increased coverage in the N-term and C-term regions. (C) Genotyping of the BC7-A spores for *HZY1* N-terminal expansion, short allele (∼580 bp) and long allele (∼2415 bp) and the correlation of long allele to the haze-positive phenotype. (D) Alignment of S288C, wine strain and OYL-011 *HZY1* alleles. (D) Schematic representation of the intragenic repeats in *HZY1* for the S288C, wine strain and OYL-011 alleles. Legend indicates identified repeat motifs along with candidate secretory and GPI anchor sequences.

We hypothesized that if the repeat expansions in OYL-011 *HZY1* allele result in the haze phenotype, then other haze-positive brewing strains may also have repeat expansions in *HZY1*. To look for potential expansions, we obtained long-read nanopore sequencing data for a collection of strains and used three highly conserved sequences upstream (tag1 ChrIX:25,878-26,392), in the central domain (tag2 ChrIX:24,727-25,335) and downstream (tag3 ChrIX:22,324-22,442) of the *HZY1* gene to extract reads containing the full N-term and C-term sequences (Figure 4A). Due to many brewing strains having heterozygous tetraploid genomes and often aneuploidies, we collected the sequence lengths from tag1-tag2 (N-term) and tag2-tag3 (C-term) to capture all potential *HZY1* alleles in each strain. The sequence lengths were then plotted using violin plots to look at both the size and distribution of the N-terminus (Figure 4B) and C-terminus (Figure 4C) in the various brewing strains. Several strains with long N-term regions had previously been identified as haze-positive (OYL-011, OYL-017, OYL-032, OYL-045) in our initial screening. Three haze-neutral strains (OYL-004, OYL-024, OYL-052) were identified to have low frequency alleles with N-term expansions (Figure 4B). We predict that copy number and expression of *HZY1* may be partly responsible for the degree of haze phenotypes observed. Also, several haze-positive strains did not have long *HZY1* alleles (OYL-061, OYL-043, OYL-015, OYL-021) and therefore it is possible that additional genes are promoting haze in these strains (Figure 4B). The C-term expansion exhibited smaller variation with the majority of expansions falling within a 500 bp distribution (Figure 4C). There was not an observed trend between length of the C-term expansions and haze, though this does not rule out the possibility of a combination of N-term and C-term expansions involved in haze. Due to the length of the reads and the size of the *HZY1* expansions, evaluation of the combinations of N-term and C-term expansions was not possible. Interestingly, OYL-001, OYL-009, OYL-077 were identified to have N-term expansions in *HZY1* and when assayed for haze were found to be among the most haze-positive strains, suggesting the N-term length of *HZY1* is partially predictive of a haze-positive phenotype (Figure 4D).

**Figure 4.**
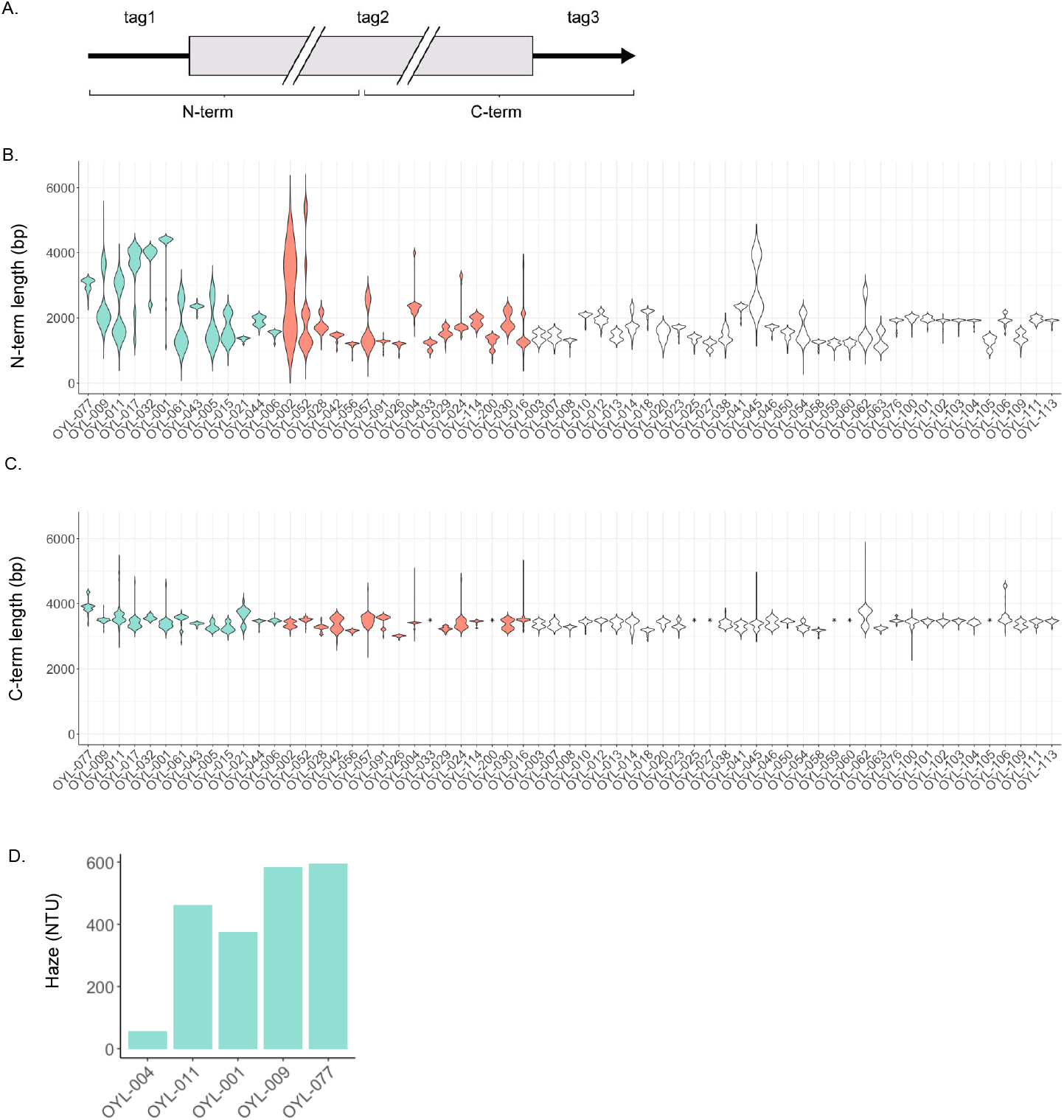
N-terminal and C-terminal expansions in *HZY1* in a collection of brewing strains. (A) Schematic of sequence tags used to extract reads from long read sequencing datasets and determine lengths of N-terminus (tag1 and tag2) and C-terminus (tag2 and tag3) in *HZY1* alleles. (B) Violin plots indicating the size and distribution of the N-terminus (region between tag1 and tag2) in long reads obtained from various brewing strains. haze-positive are indicated as green and haze-neutral as red. Strains uncharacterized for haze phenotype are white. (C)) Violin plots indicating the size and distribution of the C-terminus (region between tag2 and tag3) in long reads obtained from various brewing strains. haze-positive are indicated as green and haze-neutral as red. Strains uncharacterized for haze phenotype are white. (D) Haze phenotype of strains identified to have expanded *HZY1* N-terminus.

With a strong correlation between repeat expansions in *HZY1* to the haze phenotype, we next disrupted *HZY1* using CRISPR/Cas9 in several of the most haze-positive strains (OYL-011, OYL-009, OYL-077) as well as in one of the haze-neutral strains (OYL-004). The disruption of each allele was confirmed by PCR with complete loss of product for *HZY1* N-terminus and gain of product indicating *HZY1* disruption (Figure 5A). Each of the resulting *hzy1Δ* strains showed a substantial decrease in haze (Figure 5B). Even the OYL-004 haze-neutral strain showed reduced haze with *HZY1* disruption. This decrease in haze confirmed *HZY1* was necessary for haze formation in these strains.

**Figure 5.**
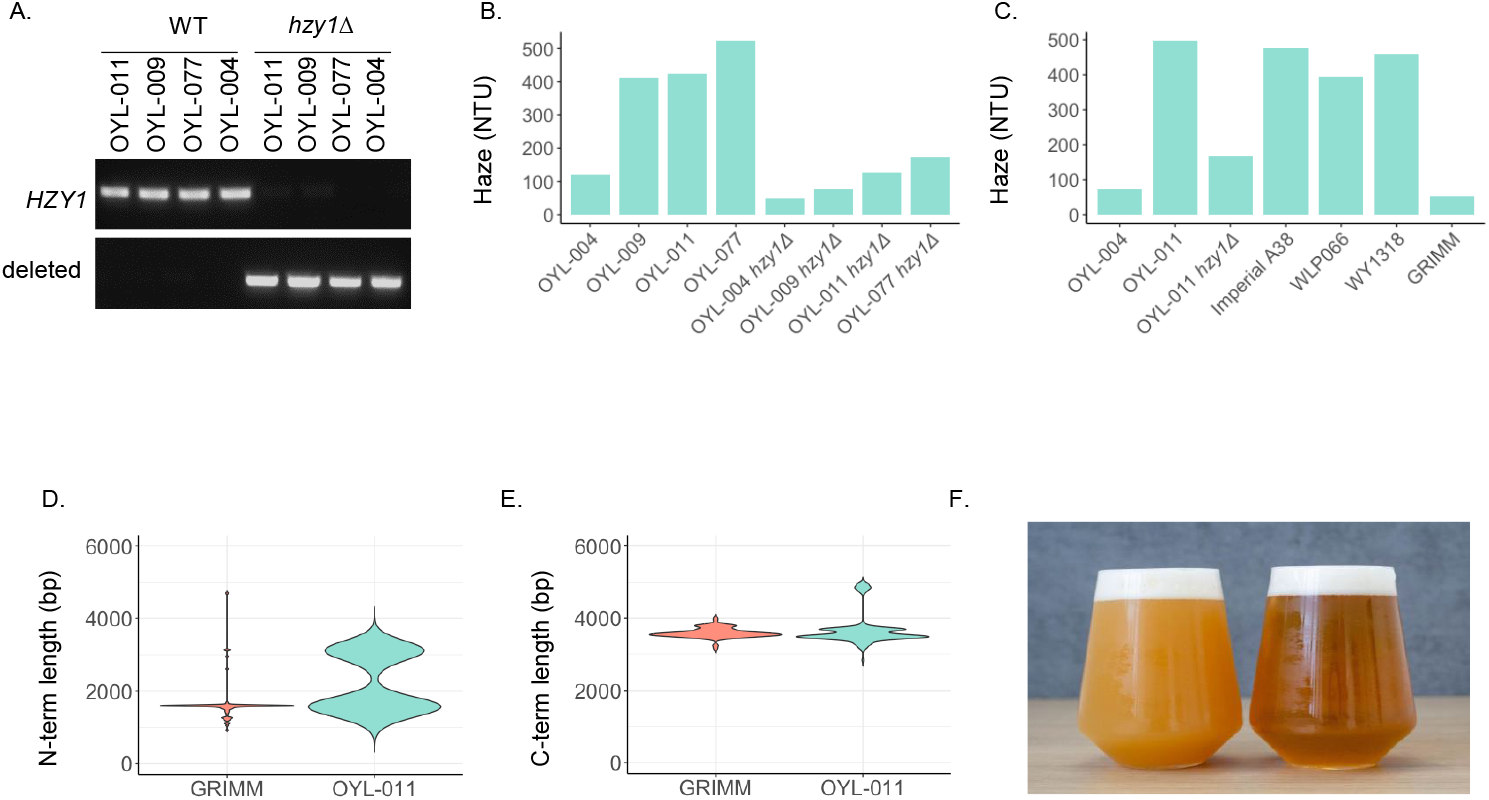
*HZY1* is necessary for the formation of dry hop-dependent haze. (A) PCR confirmation of the full CRISPR/Cas9 disruption of *HZY1* gene in all *hzy1*Δ strains. (B) The resulting haze phenotype of *hzy1Δ* strains. (C) Haze phenotypes of various commercial sources for OYL-011-equivalent strains and a generation 300 isolate from Grimm Artisanal Ales (GRIMM). (D) Distribution of *HZY1* N-term length in GRIMM and OYL-011. (E) Distribution of *HZY1* C-term length in GRIMM and OYL-011. (F) Typical IPA recipe fermented with OYL-011 (left) and OYL-011 *hzy1*Δ (right).

While this study was underway, we were in touch with the brewery owners at Grimm Artisanal Ales who had been having difficulty obtaining stable haze using their house culture (GRIMM), which was a derivative of OYL-011 from another commercial yeast supplier. A sample of the GRIMM strain was tested and found to be haze-neutral. The brewery had maintained their house culture for 300 generations by top cropping yeast in active fermentation and utilizing it for successive fermentations. We then assayed several of the other yeast supplier equivalent strains for OYL-011 and found all but the GRIMM isolate to be haze-positive (Figure 6C). We obtained long-read sequencing on the GRIMM isolate to see if *HZY1* had undergone genetic changes that might result in the loss of the haze phenotype. Surprisingly, the GRIMM strain differed from the OYL-011 strain in that it no longer had expansions in the N-term or C-term of *HZY1* (Figure 5D/5E). The loss of a long allele in the non-hazy GRIMM isolate suggests that expansion and contraction (or possibly *HZY1* disruption) can result in the gain and loss of the haze phenotype in a brewery setting and provides further evidence for the role of expanded *HZY1* alleles in promoting haze.

The typical beer consumer is heavily influenced by the appearance of beer and haze is often associated with either an increase or decrease in hedonic liking. We set out to determine whether this is strictly based on appearance and associations to beer styles, or if haze also has an impact on the taste and aroma of the beer. A standard IPA recipe was brewed and split into two fermentations, one with OYL-011 and one with OYL-011 *hzy1Δ*. The haze in the resulting beers measured 428 NTUs and 40 NTUs, respectively. This difference was very visually striking and would present very differently to the beer consumer (Figure 5F). A tetrad sensory test was performed where the beer samples were kept covered and in opaque cups to prevent panelists from determining which was hazy or not hazy. Only one out of eleven panelists was able to identify the correct pairing, indicating that the aroma, mouthfeel and taste were not statistically different between the hazy and non-hazy beers.

## Discussion

This study investigated the genetics of beer haze in *Saccharomyces cerevisiae*. Much of the haze research in beer to date has focused on malt and hop contributions but a role for yeast has remained largely unknown. From our initial experiments characterizing dry hop-dependent haze in a collection of commercial brewing strains, it became apparent that brewing strains exhibit a wide range of haze phenotypes (referred to as haze-positive and haze-neutral). Employing a classic genetic backcrossing approach, we identified YIL169C, which we have termed *HZY1*, as a novel haze gene. Through *HZY1* knockout experiments and the correlation between the expansion of *HZY1* intragenic repeats and haze-positive phenotype amongst brewing strains, we were able to provide further evidence for *HZY1* promoting the haze phenotype. Together our results provide the first evidence that the *S. cerevisiae* gene, *HZY1*, has a critical role in promoting haze in dry-hopped beer styles.

The *HZY1* gene encodes a mannoprotein with a putative GPI anchor and includes intragenic repeats in both the N-terminal serine-rich and C-terminal serine/threonine-rich domains. *HZY1* intragenic repeats possess a high degree of genomic plasticity with massive expansions resulting in a range of haze phenotypes observed amongst brewing strains. The majority of whole genome sequencing studies with brewing strains have employed short read sequencing technologies ^18^. Though short read data provides a highly accurate coverage of the genome, complex repetitive sequence cannot be resolved and thus *HZY1* has largely remained uncharacterized. *HZY1* also is located in the subtelomeric region of chromosome IX, and these regions are often genomic hotspots for mitotic recombination and interchromosomal reshuffling, which partially can help to explain the degree of genomic and phenotypic variability that we have uncovered in this study. In addition, the complex genetics of domesticated *S. cerevisiae* brewing strains including heterozygosity, polyploidy, and chromosomal rearrangements might also suggest that domesticated yeast have a higher degree of evolution in the *HZY1* gene ^19^. In future studies, long-read sequencing approaches can help to resolve and phase different *HZY1* alleles across wild and domesticated strains. A better understanding of allele types and frequencies will help to further elucidate this complex haze trait.

Flocculation and maltose utilization traits exhibit a high degree of biodiversity within brewing strain phenotypes and genotypes. There are clear selective pressures for these two phenotypes as brewers often harvest from the bottom of the fermentation vessel and use wort primarily composed of maltose. However, the selective pressure for haze and any possible fitness advantage to *HZY1* intragenic expansions remains unknown. The expansion of these serine and threonine-rich regions in Hzy1 could lead to heavy glycosylation which could change the cell surface composition, possibly altering cellular adhesion and responses to environmental stress. Among *S. cerevisiae* genes with long intragenic tandem repeats, 75% encode for cell-surface proteins. Many of these genes also exhibit expansions and contractions in the intragenic repeats, including *FLO1, FLO5, FLO9, FLO11* and *HPF1* and impact cell adherence and buoyancy ^17,20^. We have not found an association between haze-positive and flocculation phenotypes within brewing strains, but it is possible that the *HZY1* repeat expansions were selected for due to their effect on cell buoyancy. Interestingly, the lager strains analyzed in this study exhibited little diversity in *HZY1* relative to ale strains and some of the hazier ale strains have been historically top cropped. If *HZY1* expansion leads to more buoyant cells, then these cells would be preferentially selected through top cropping, similar to the flocculant cells being selected through bottom cropping.

Our study provides the first evidence for the involvement of *S. cerevisiae* and the gene, *HZY1*, in the promotion of dry hop-dependent haze. The study will likely fuel future research investigating additional genes involved in haze, as well as biochemical studies to determine the composition and the biophysical properties of beer haze. Ultimately, the knowledge shared within will leave brewers with a deeper understanding of yeast-derived haze and the strains and methods that can be used to perfect their hazy (or non-hazy) brands.

## Data Access

Long-read sequencing data for *HZY1* can be provided upon request.

## Acknowledgements

We thank all our team members at Omega Yeast Labs with special thanks to Allison Lange, Chris Bernardo and Nate Morton for many discussions, and Dexter Stephens for his illustrations. We thank Joe Grimm, Lauren Carter Grimm, Justin Knoll and the Grimm Artisanal Ales team for their curiosity and for sharing their house culture. We also thank Felix Seifert for bioinformatics support. The strain sequencing and analysis was supported in-part by the USDA National Institute of Food and Agriculture Multistate Hatch funds (NC1023).

---

**Supplemental Figure 1.**
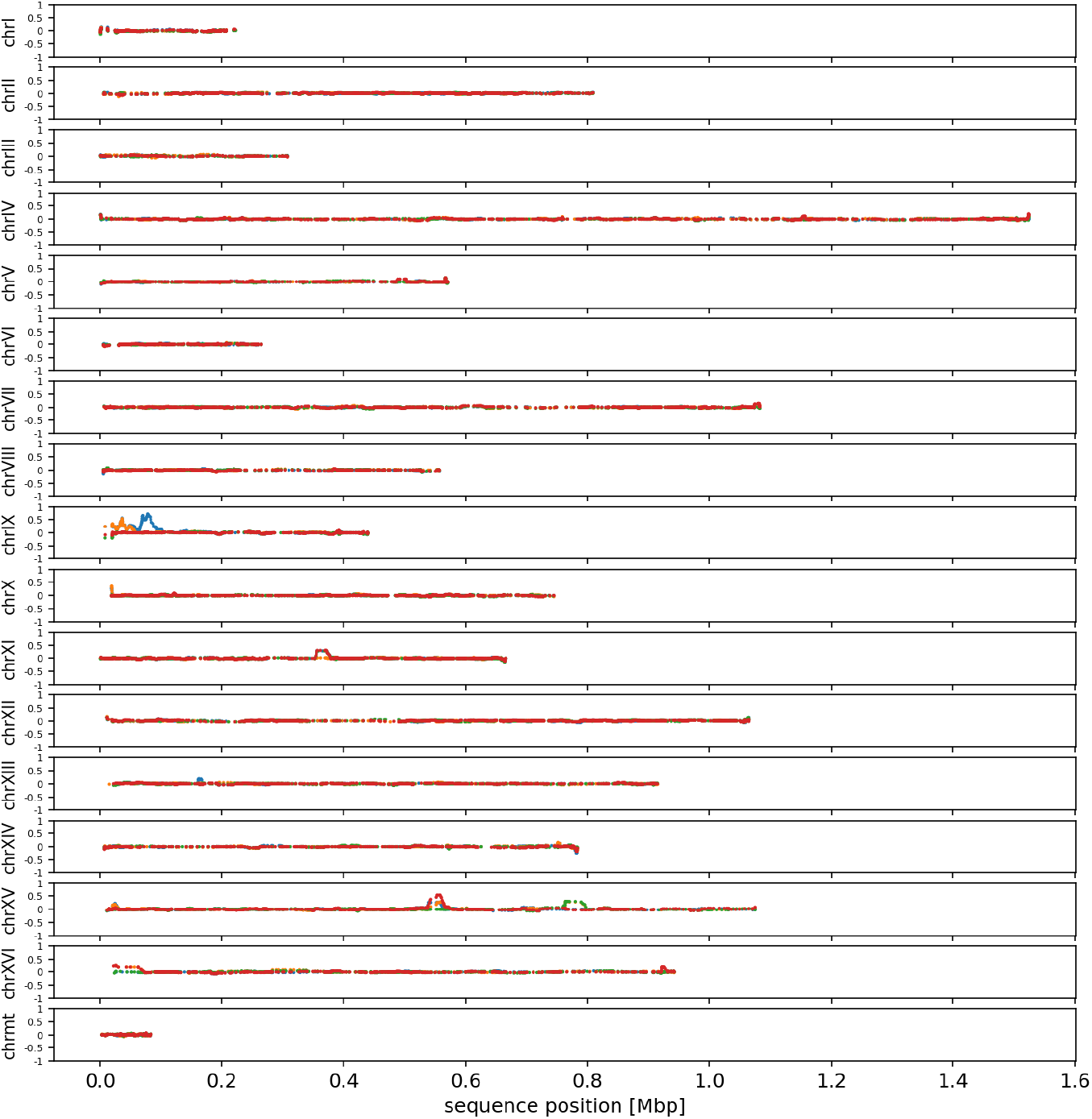
The ratio of variants found in the BC7 isolates relative to the parent wine strain (y-axis) are plotted with a sliding window of 50 bp along the chromosome coordinates of the reference genome S288C (x-axis). BC7-A in blue, BC7-B in orange, BC7-C in green and BC7-D in red.

